# DNA from dried blood spots yields high quality sequences for exome analysis

**DOI:** 10.1101/2020.05.19.105304

**Authors:** Uma Sunderam, Aashish N. Adhikari, Kunal Kundu, Jennifer M. Puck, Robert Currier, Pui-Yan Kwok, Steven E. Brenner, Rajgopal Srinivasan

**Author notes:** Equal contribution from these authors (Steven E. Brenner, Rajgopal Srinivasan).

## Abstract

**Background:** DNA sequencing of archived dried blood spots (DBS) collected by newborn screening programs constitutes a potential health resource to study newborn disorders and understand genotype-phenotype relationships. However, its essential to verify that sequencing reads from DBS derived DNA are suitable for variant discovery.

**Results:** We explored 16 metrics to comprehensively assess the quality of sequencing reads from 180 DBS and 35 whole blood (WB) samples. These metrics were used to assess a) mapping of reads to the reference genome, b) degree of DNA damage, and c) variant calling. Reads from both sets mapped with similar efficiencies, had similar overall DNA damage rates, measured by the mismatch rate with the reference genome, and produced variant calls sets with similar Transition-Transversion ratios. While evaluating single nucleotide changes that may have arisen from DNA damage, we observed that the A>T and T>A changes were more frequent in DNA from DBS than from WB. However, this did not affect the accuracy of variant calling, with DBS samples yielding a comparable count of high quality SNVs and indels in samples with at least 50x coverage.

**Conclusions:** Overall, DBS DNA provided exome sequencing data of sufficient quality for clinical interpretation.

## Background

Several countries routinely collect neonatal dried blood spot (DBS) samples as part of their newborn screening (NBS) programs. The blood is typically collected onto filter cards from the heel of newborns, air dried, and subsequently used in screening for diseases [1]. Archived collections of DBS samples stored desiccated at −20° C provide a rich source of material for scientific inquiry aimed at establishing genotype to phenotype correlations.

In collaboration with the California Department of Public Health (CDPH), we engaged in a project to assess the potential applications of whole exome sequencing (WES) in a NBS program to discover rare disorders in clinically asymptomatic children. Our initial pilot study used whole WES to identify metabolic disorders that were already being screened for in NBS programs using tandem mass spectrometry (MS-MS).

To identify genetic variants associated with various diseases from the DNA extracted from DBS samples stored for several years, one has to first evaluate whether the quality of the extracted DNA sequences can provide exomes comparable to that derived from whole blood (WB) DNA. This is especially important for studies of rare Mendelian disorders, for which analysis involves locating a single causative genetic variant across the whole exome. Sufficient DNA needs to be extracted to provide adequate coverage of regions of interest, as each DBS sample contains only about 50 µl of blood [2]. Furthermore, any DNA damage to the DBS samples could potentially impact variant calling and needs to be examined.

A previous study compared pairwise variant concordance rates as a similarity measure between matched DBS and whole blood samples [3], concluding that whole genome amplified DNA from DBS gave comparable WES to whole blood (WB). The cohort size of 22 was much smaller than ours, and the focus was on variant analyses; other metrics reported were minimal. Another study successfully used WES from DBS for identification of important genes in a specific disorder, bronchopulmonary dysplasia, using case and control data [4]. A retrospective clinical study developed a targeted gene panel and variant processing pipeline to demonstrate utility for NBS [5]. However, these studies did not extensively compare the quality of WES from DBS versus WB and had smaller cohort sizes than our study.

Our study evaluated the exome quality of DNA from DBS for use in screening for monogenic autosomal recessive Mendelian disorders in which two rare variants in a single gene must be found to explain the disease. Hence, the individual variant level quality needs to be good for every sample, since missing a variant because of low quality data can lead to misdiagnosis of a life-threatening condition. In the absence of matched samples, we focused on an extensive set of metrics that span the entirety of the process of variant calling starting from raw sequence reads. We compared metrics from two datasets: i) 180 DBS samples (DBS_2006) archived between the years 2006 and 2013; and ii) 35 WB samples collected, extracted and sequenced between 2013 and 2014. In addition, we evaluated a set of 8 anonymous DBS samples (DBS_1980) archived between 1980 and 1982 for the same set of parameters (Additional file1; DBS_1980 metrics section).

## Results

To assess the quality of the sequencing we evaluated three categories of quality metrics (Table 1) across both data sets:

- Read mapping to the reference sequence (Mapping Quality Metrics)
- DNA damage in reads (DNA Damage Metrics)
- Called variants (Variant Quality Metrics)

**Table 1.**
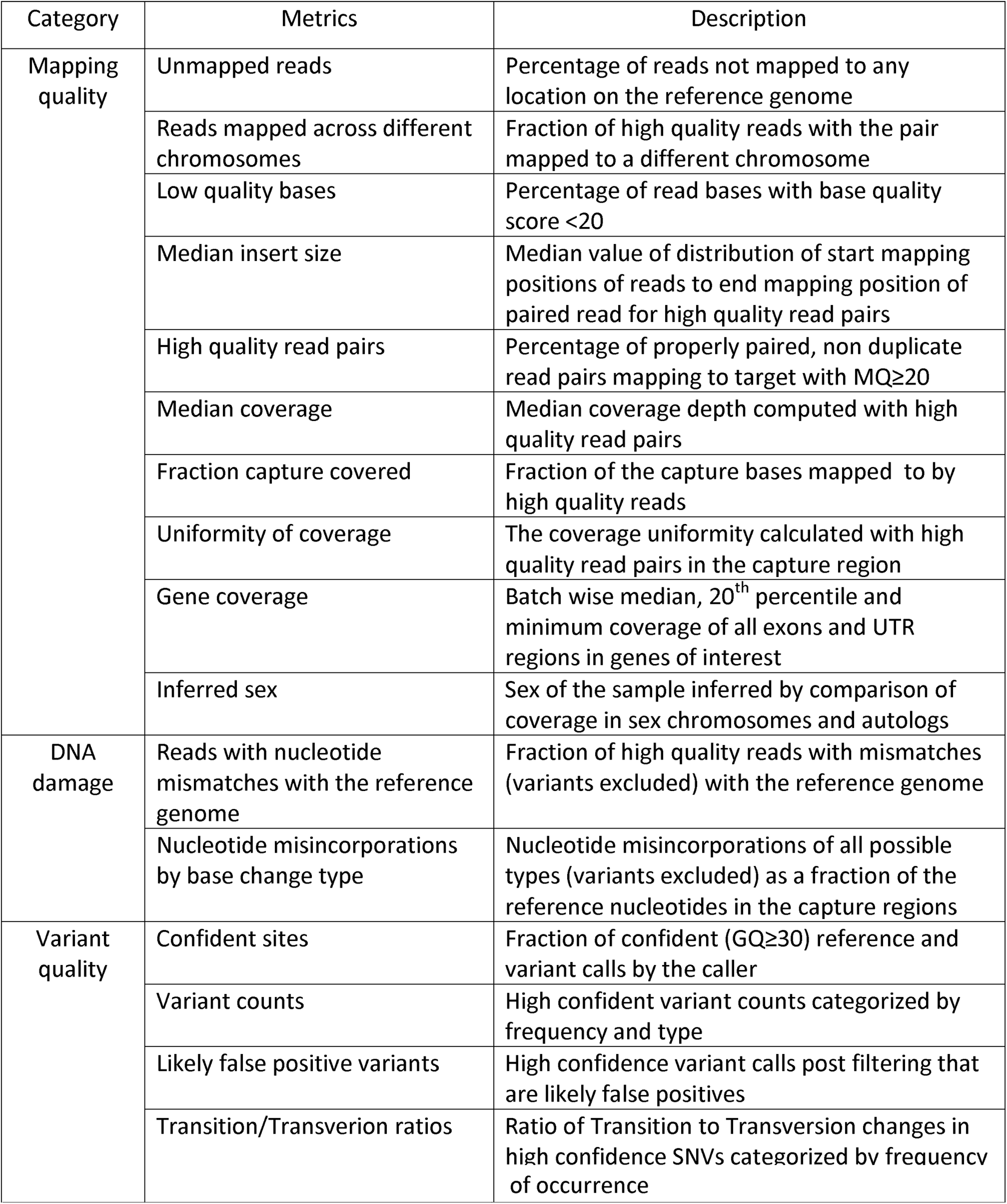
Metrics for comparison of DBS with WB DNA sequence data.

### Mapping Quality Metrics

Successful variant calling requires raw reads to be mapped to the reference genome with high confidence and in sufficient numbers across all regions. We scrutinized parameters at nucleotide level and read mapping level. DBS_2006 data were in expected ranges and similar to WB values for unmapped reads; low quality bases; read pairs mapped across chromosomes; median insert sizes; and on-target, high quality read pairs (Additional file1; Mapping quality metrics section).

### Capture Coverage

Adequate coverage across the exome capture region is an indicator that sufficient quantity of high quality DNA was extracted to make confident variant calls. Coverage different from that expected from the library preparation and sequencing can indicate poor mapping. We compared the median depth of coverage across the capture region as well as fraction of the capture region covered at various depth cutoffs ranging from 1x to 30x. We used high quality read pairs mapped to the capture region (Additional file1; High quality reads pairs section) for all coverage-related metrics unless otherwise specified. The WB set had a higher median coverage for most of the samples (Table 2; Additional file1, Figure S2A). The median coverage correlated strongly with the total number of reads for a sample (correlation coefficients-WB: 0.98, DBS_2006: 0.96; Additional file1, Figure S2C). The WB samples in our study had been sequenced to a greater depth and hence had on average higher number of reads than the DBS samples (Additional File1; Figure S1A).

**Table 2.** Coverage related metrics for WB and DBS_2006 datasets.

| Metric | Median |  | 1<sup>st</sup> quartile |  | 3<sup>rd</sup> quartile |  | p-value |
| --- | --- | --- | --- | --- | --- | --- | --- |
|  | WB | DBS_2006 | WB | DBS_2006 | WB | DBS_2006 |  |
| Median Coverage | 84 | 53 | 53.5 | 48 | 96.5 | 59.25 | 3E-06 |
| Fraction covered at 1x | 0.98111 | 0.98176 | 0.97955 | 0.980925 | 0.98228 | 0.9824725 | 0.01 |
| Fraction covered at 5x | 0.96707 | 0.969215 | 0.96516 | 0.9673875 | 0.97392 | 0.9705675 | 0.7 |
| Fraction covered at 10x | 0.95598 | 0.95702 | 0.954875 | 0.9536975 | 0.964 | 0.95949 | 0.009 |
| Fraction covered at 15x | 0.94721 | 0.943275 | 0.945935 | 0.9363 | 0.951705 | 0.9484125 | 0.002 |
| Fraction covered at 20x | 0.93739 | 0.92207 | 0.930955 | 0.9077575 | 0.94022 | 0.93356 | 0.005 |
| Fraction covered at 30x | 0.90245 | 0.841045 | 0.8606 | 0.805715 | 0.92105 | 0.878385 | 0.0009 |
| Uniformity of coverage | 34.6 | 31.9 | 32.3 | 31.1 | 35.8 | 32.8 | 7E-05 |
| IQR/median coverage | 0.72727 | 0.66667 | 0.67424 | 0.64706 | 0.75192 | 0.68421 | 0.0002 |

The DBS_2006 exhibited marginally better breadth of coverage compared to the WB set with fraction of the capture intervals covered at 1x being better (Table1; Additional file1, Figure S2B). The fraction of capture regions covered at 5x, 10x and 15x were comparable in both sets. Since the DBS_2006 was sequenced to a lower depth compared to the WB, comparisons could not be done beyond 30x.

### Uniformity of Coverage

Exomes are characterized by non-uniform regions of coverage compared to whole genome sequencing data [6]. Lower coverage at a particular locus can cause true variants to be missed. The uniformity of coverage was evaluated as follows:

For each position in the capture, we defined deviation from mean coverage as

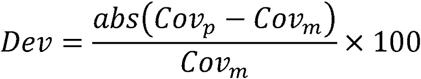

where Cov_m_ is mean coverage across the capture region and Cov_p_ is coverage depth at position p. Hence, the fraction of positions with coverage within i% of Cov_m_ is

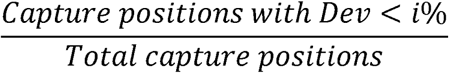

where i = 10 to 90% deviation from mean, in steps of 10.

We plotted the deviation from the mean coverage % and the corresponding fraction of base positions (median data across each set) for WB and DBS_2006. A steeper gradient implies more uniform coverage in the sample (Figure 1A), since for a given distance from the mean, this would translate to a larger percentage of bases within that coverage. We interpolated the data to calculate the uniformity of coverage as the maximum deviation from the mean coverage in 50% of the base positions and compared the value for the two data sets. This was lower in DBS_2006 compared to the WB samples (Table1; Figure 1B). Five WB samples appeared to deviate significantly from the mean compared to the others.

**Figure 1:**
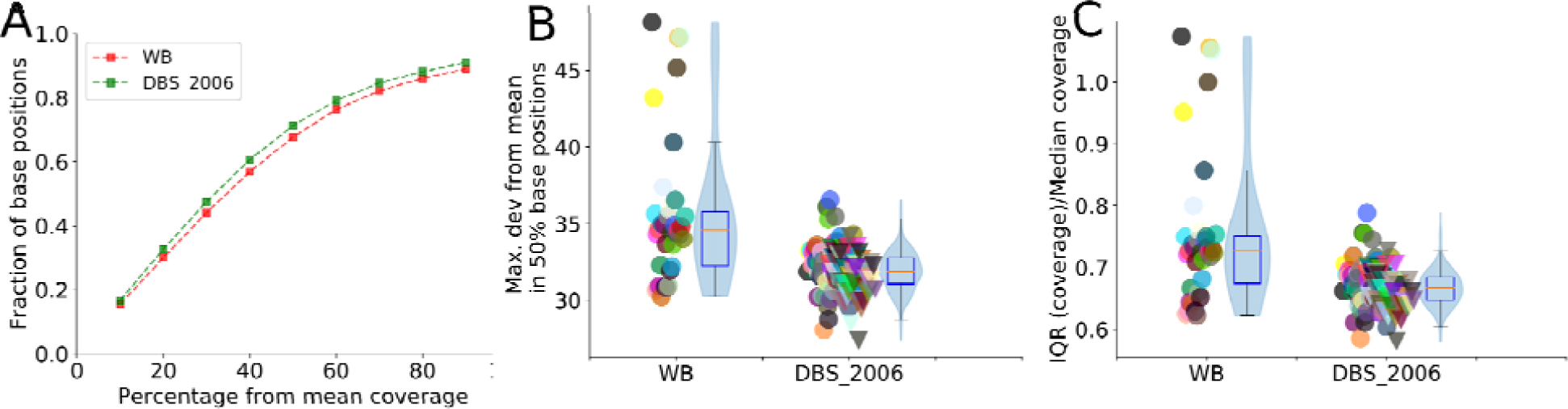
Uniformity of coverage in WB and DBS_2006. A: Fraction of total positions in capture (median value for each data set) is plotted on y-axis and maximum deviation from mean of that fraction on the x-axis. A) Red: WB, B) B) Black: DBS_2006. The DBS_2006 set, showed more uniformity of coverage, with the curve showing a steeper gradient. B: Maximum deviation from mean in 50% capture sites in WB and DBS_2006 data sets. Deviation from mean was lower in the DBS_2006 compared to the whole blood dataset. C: Inter-quartile range coverage/Median coverage in WB and DBS_2006 data sets. This was lower in most of the DBS_2006 samples compared to the WB dataset. In B and C, Individual sample values are plotted and adjacent box plots display the median (red) and interquartile ranges for the dataset, whiskers extend to the last data point within 1.5 times the interquartile range. Violin plots superimposed on the box plots show the data density and mean value (blue).

Another measure of uniformity of coverage is

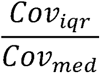

where Cov_iqr_ is the inter-quartile coverage depth of all base positions in the sample and Cov_med_ is the median coverage. The lower this value, the smaller the deviation from the median coverage and hence the more uniform the coverage. The inter-quartile coverage depth divided by median coverage also showed a similar trend with the DBS_2006 set having lower values compared to the WB data (Table 2; Figure 1C).

### Gene Coverage

We examined base-wise coverage for exons from a set of 78 genes known to be associated with Mendelian inborn errors of metabolism as part of a study that examined the role of exome sequencing in new born screening [7]. The median coverage across all samples, 20th percentile coverage and lowest coverage for all positions were plotted (Figure 2). Genes that were well covered in the WB set were also well covered in the DBS_2006 set. Data for two such genes, ACADM and CPT1A, are shown in the figure. Similarly, exons in some genes were observed to have low coverage in both WB and DBS_2006. Data for two such genes, BCKDHB and MLYCD, are shown in Figure 2, and are consistent with an artifact of the capture.

**Figure 2:**
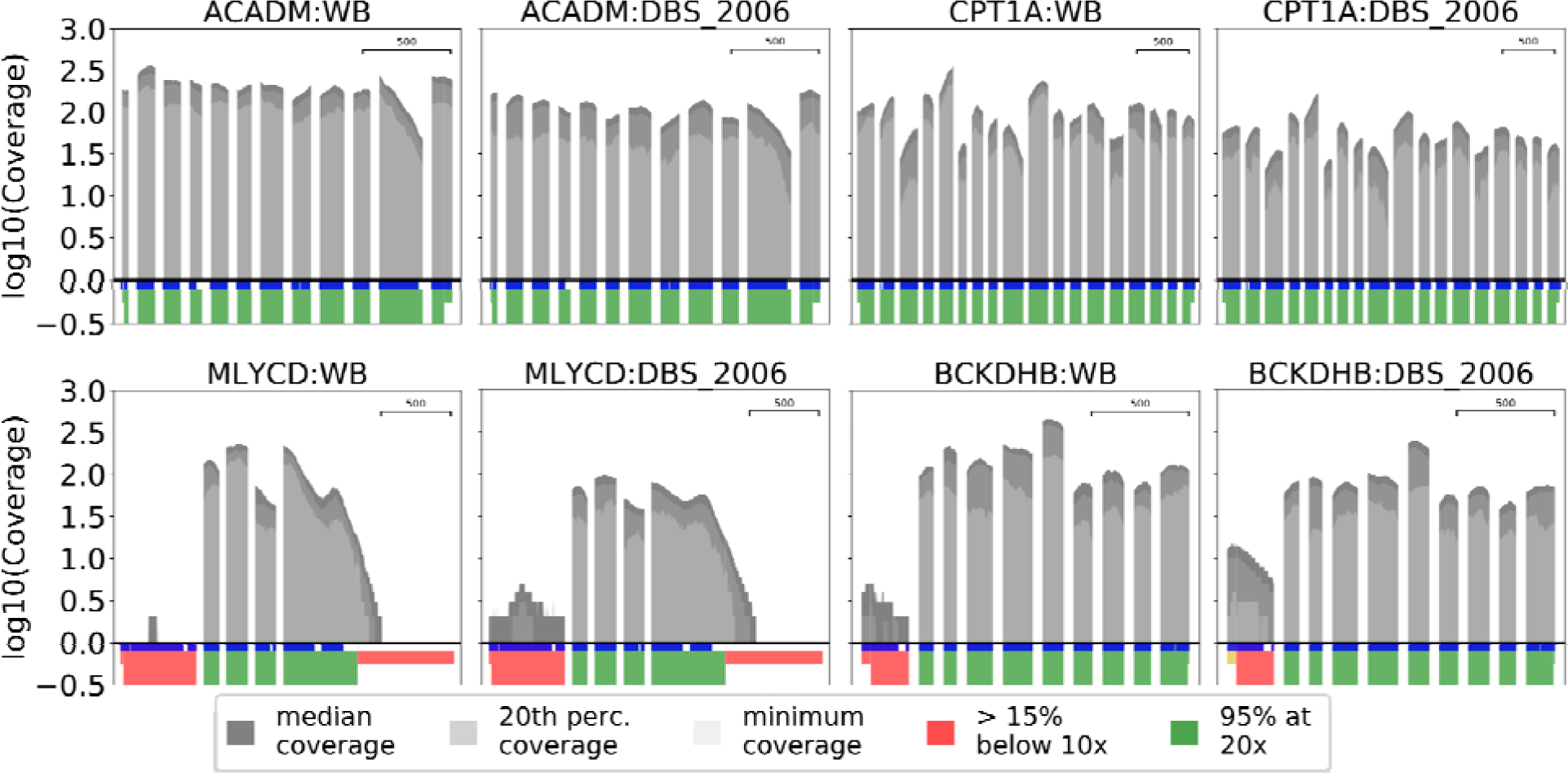
Median coverage for whole blood (WB) and recent dried blood spot (DBS_2006) samples. Row 1: Coverage plots for ACADM and CPT1A genes. Row 2: Coverage for CPT1A and BCKDH genes. Plot of log(base 10) of the median, 20 ^th^ percentile and minimum coverage for each coding exon across all samples for a given sample set. Dark grey: Median coverage, medium grey: 20^th^ percentile coverage, light grey: minimum coverage at that position respectively. Coverage quality of each exon is indicated by colored blocks beneath the exon. Coverage quality of each exon is indicated by colored blocks beneath the coverage plot. Red: Greater than 15% of exon has less than 10x median coverage; green: 95% of the exon has minimum 20x coverage. UTRs that are part of the coding exons have a smaller indicator thickness. Regions of the exon that overlap with the capture array are indicated in blue just below the coverage plot. Exon scale in bases is shown in each plot. The well covered and poorly covered genes showed a similar trend in all 3 datasets. ACADM and CPT1A are examples of well covered genes and BCKDH and MLYCD have poorly covered exons. The coverage trends were similar at exon level as well.

### Inferred Sex

The recorded sex of the individual submitting the sample was inferred by a chromosome coverage. Inconsistency of sex inferred from DNA versus sex recorded at the time of sample collection is an indicator that the DNA is poorly mapped and of insufficient quality.

We used chromosome wise coverage information across the capture intervals to infer the sex of the samples. The coverage metric for a chromosome i is defined as

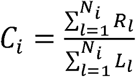

where R_l_ = Number of reads overlapped with the capture interval l in chromosome i, L = Length of capture interval, N_i_ = Number of capture intervals in I for the sex chromosomes and 2 chromosomes with similar length of capture intervals, 4 and 15. Only non-PAR regions of the X and Y chromosomes were considered.

The distance calculations were

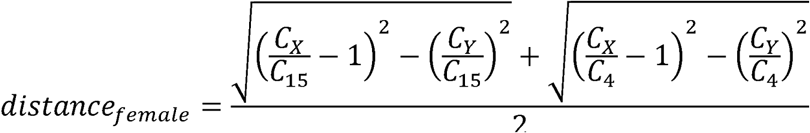

and

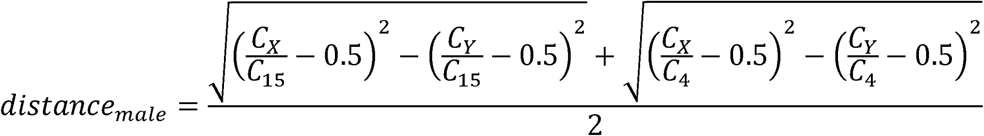

where C_X_, C_Y_, C_4_ and C_15_ are the coverage coefficients for the respective chromosomes. The inferred sex for a given sample is assigned according to the minimum value of distance_male_ and distance_female_. The inferred sex in all cases agreed with the reported sex for the WB and DBS_2006 data sets.

### DNA damage metrics

DNA damage involving chemical modification of the template nucleotides can lead to erroneous incorporations of nucleotides during the synthesis of the complementary DNA [8]. These base changes with respect to the reference genome are not true SNVs and impact the quality of the final call set, leading to a higher false positive variant call rate. Furthermore, if there is DNA damage on the reads with true variants, the confidence of the true variants is decreased as the mapping quality could be lower, thus leading to true variants being flagged as potential false positives during variant filtering. We evaluated the following metrics to assess degree of DNA damage in the DBS sets and the WB samples. Both metrics were calculated using high quality read pairs (Additional file1; High quality reads pairs section).

### Reads with nucleotide mismatches with respect to the reference genome

DNA stored for long periods could potentially accumulate various base alterations. A previous study estimated DNA damage by comparing the number of mismatches in the aligned reads between germline and FFPE (formalin-fixed paraffin embedded) tumor samples and found that the proportion of reads with ≥1 mismatched base were greater in the FFPE tumor samples [9]. Following this approach, we calculated the fraction of mismatch bases as

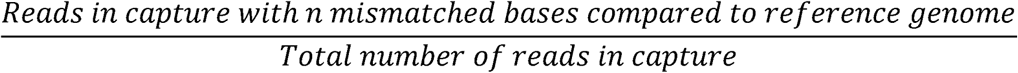

where n = 0, 1, 2 and ≥3 mismatches. We excluded variants called with high confidence that overlapped with the mismatch positions to ensure true variants were not counted as mismatches.

Both datasets showed similar fractions of mismatch reads (Table 3). In both data sets, >80% of reads had no mismatches to the reference genome in most samples (WB, 88.6%; DBS_2006, 99.4%, as shown in Figure 3). Reads with ≥3 mismatches were slightly more frequent in the WB set compared to the DBS_2006 data (Table 3).

**Table 3.** Mismatch bases with the reference genome for WB and DBS_2006 datasets.

| Metric | Median |  | 1<sup>st</sup> quartile |  | 3<sup>rd</sup> quartile |  | p-value |
| --- | --- | --- | --- | --- | --- | --- | --- |
|  | WB | DBS_2006 | WB | DBS_2006 | WB | DBS_2006 |  |
| 0 mismatches | 0.90106 | 0.89756 | 0.88022 | 0.89152 | 0.90847 | 0.90478 | 0.1 |
| 1 mismatch | 0.07291 | 0.07792 | 0.0654 | 0.07145 | 0.07879 | 0.08276 | 0.2 |
| 2 mismatches | 0.01252 | 0.01283 | 0.01215 | 0.01187 | 0.01821 | 0.0142 | 0.2 |
| ≥3 mismatches | 0.01347 | 0.01163 | 0.01255 | 0.01003 | 0.01812 | 0.01487 | 0.01 |

**Figure 3:**
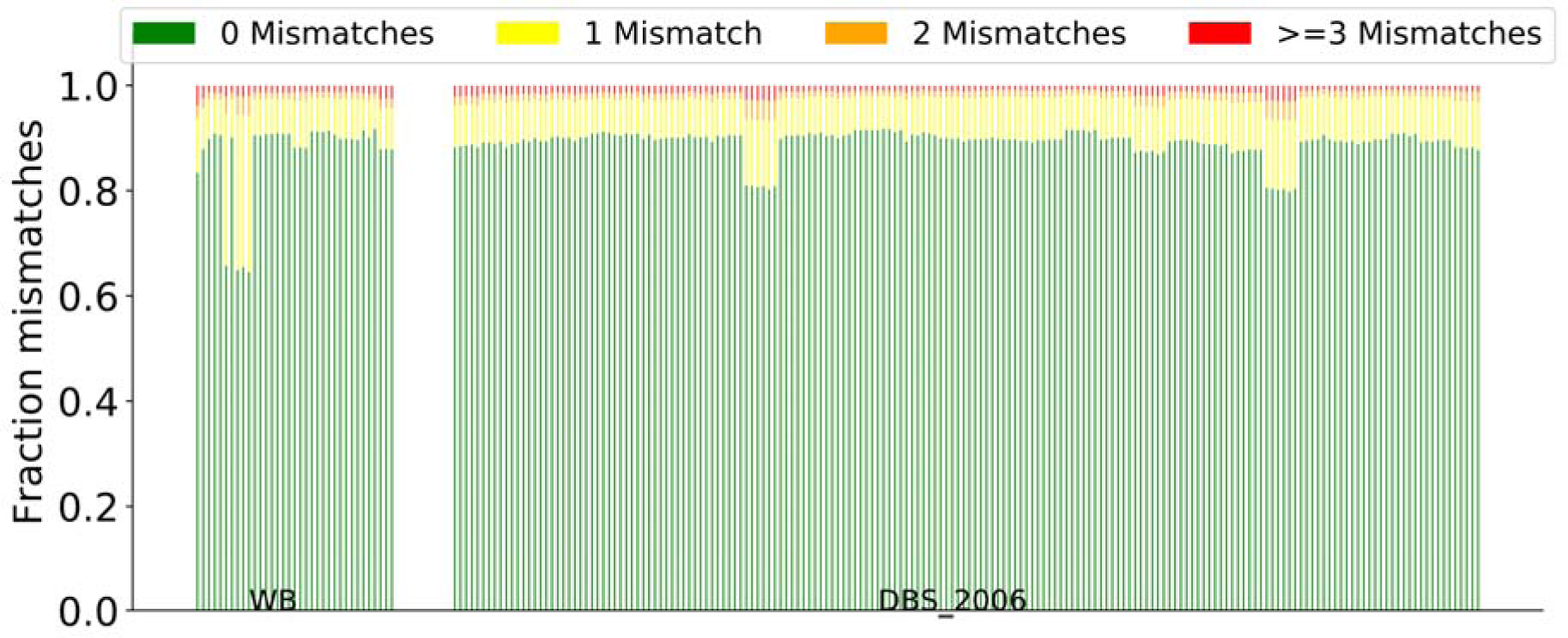
Fraction of reads with 0, 1, 2 and 3 or more mismatches with the reference. Both WB and DBS_2006 datasets had similar ranges of fraction reads with 0, 1, 2, and 3 or more mismatches at positions not called variants by GATK Haplotype Caller.

### Nucleotide misincorporations by base change type (NMBC)

Relative frequencies of different kinds of non-variant base changes have previously been studied in FFPE in comparison with frozen tissue samples as an estimate of DNA damage [10]. C>T and G>A transitions were found to be significantly more frequent than other changes in the FFPE set. The largest C>T difference was seen in the dinucleotide CG>TG transitions. These changes could potentially result in erroneous variants being discovered by the variant caller.

We calculated the frequency of the nucleotide misincorporations by base type for a sample as

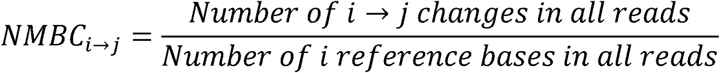

where I,j = A, C, T, G and i≠j. For instance, the fraction of A>C changes was computed as the ratio of the number of times an A>C change was seen in the sample to the total number of A nucleotides seen. Similarly, we computed all possible base changes (A>C, A>G, A>T, C>A, C>G, C>T,….T>G). Only high quality read pairs occurring in the capture region as described previously were considered, and reads with insertions and deletions were excluded. To distinguish nucleotide misincorporations from true variants, we excluded high confidence variants from the computation (Methods; Reads and variant selection for metrics computation). We might expect to see more C>T changes in the DBS data as compared to the WB samples, given that deamination of cytosine and subsequent conversion to thymine is a recognized source of spontaneous mutation [11]. However, the C>T misincorporations were comparable among both sets. The A>T and T>A changes were higher in the DBS_2006 set compared to the WB, whereas A>G, A>C, C>G, G>C, T>C and T>G changes were higher in the WB set compared to the DBS_2006 samples (Additional file2, Table S2B; Figure 4).

**Figure 4:**
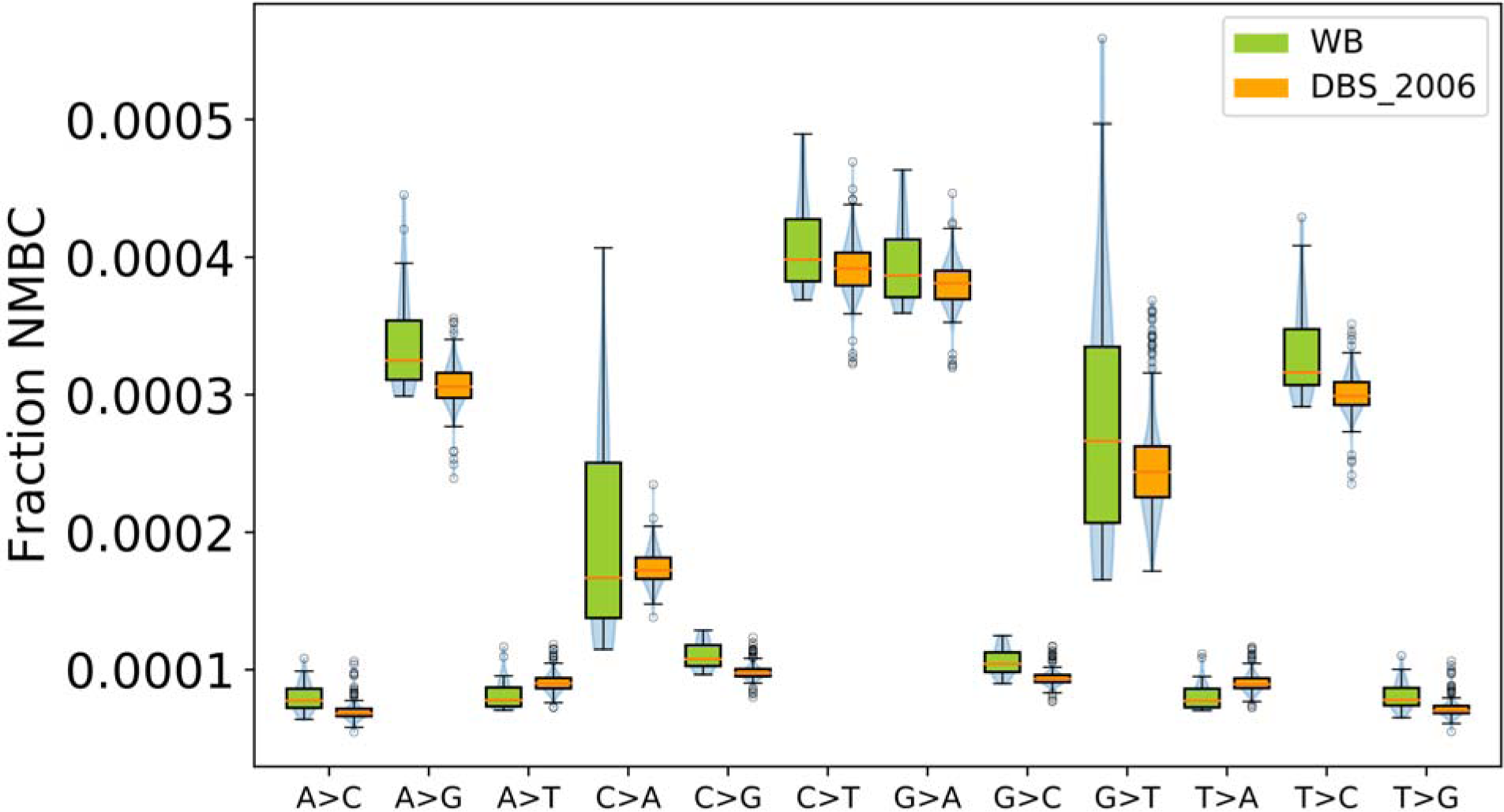
Fraction nucleotide misincorporations by base change (NMBC) for all single base changes. Box plots display the median and inter quartile ranges for the dataset, whiskers extend to the last data point within 1.5 times the interquartile range and outliers beyond this are shown as circles. Green: WB, Orange: DBS_2006. The A>T and T>A changes were slightly higher in the DBS_2006 samples compared to WB.

### Variant Calls

This set of metrics assessed the quality of the variants ultimately called by the variant caller. High quality variants (Methods; Reads and variant selection for metrics computation) were analyzed for both data sets. These were categorized into common variants, with a minor allele frequency ≥0.001, and rare variants, with a minor allele frequency <0.001 in the 1000 Genomes database (phase3) [12]. Only variants in the regions defined in the Nimblegen capture array (v3) [13] bed file were considered for the analysis.

High quality variants not present in the 1000 Genomes variant set, but present in more than a few samples in our study cohort were likely false positives. These could arise from systematic sequencing errors or through DNA damage.

### Confident Sites

We compared the reference and variant sites called confidently by the variant caller as a fraction of the capture region between the DBS_2006 and WB sets. A higher fraction of such confident calls would increase confidence in downstream variant interpretation. For each individual sample, the Genomic VCF (GVCF) files generated by the Haplotype Caller from GATK was used. All sites with genotype quality ≥30 were counted as confident and the total fraction of such sites with respect to the all the sites in the capture region was computed. The WB and DBS_2006 data had fractions of confident sites in similar ranges (Table 3; Additional file1, Figure S3A).

### Variant Counts

The count of common and rare variants were compared between the WB and DBS sets. These were further categorized Single Nucleotide Variations (SNVs) and Indels. To compare the variants, the WB, DBS_2006 and DBS_1980 samples were called together (223 samples).

The counts of high quality, common SNVs were comparable in the WB and DBS_2006 data (Table 4; Figure 5). The rare SNV frequencies were also comparable. Common indels were more frequent in the WB set, while rare indels were similar in frequency. In addition, to mitigate the influence of each data set on the other in calling variants, WB and DBS data were called separately – 35 WB samples together and 180 DBS_2006 samples together. The variants were compared in these two sets (Additional file 1; variants – additional call sets section).

**Table 4.** Variant statistics for WB vs DBS_2006 datasets.

| Metric | Median |  | 1<sup>st</sup> quartile |  | 3<sup>rd</sup> quartile |  | p-value |
| --- | --- | --- | --- | --- | --- | --- | --- |
|  | WB | DBS_2006 | WB | DBS_2006 | WB | DBS_2006 |  |
| Confident sites | 0.95161 | 0.9518 | 0.95023 | 0.94738 | 0.95851 | 0.95485 | 0.004 |
| Common SNVs | 69328 | 68905 | 68548 | 68110.3 | 70106 | 69564.3 | 0.1 |
| Rare SNVs | 835 | 795.5 | 781 | 731.8 | 1026.5 | 929 | 0.05 |
| Common Indels | 5281 | 5148 | 5232.5 | 5094.8 | 5326 | 5223.3 | 9E-05 |
| Rare Indels | 416 | 401 | 393.5 | 386.8 | 425.5 | 422.3 | 0.1 |
| LFP SNVs | 1078 | 1031.5 | 1048 | 989.8 | 1113 | 1080.3 | 0.0009 |
| LFP Indels | 1991 | 1984.5 | 1901.5 | 1939.5 | 2042.5 | 2041.3 | 0.5 |
| Ti/Tv - Common SNPs | 2.56 | 2.55 | 2.54 | 2.54 | 2.56 | 2.56 | 0.9 |
| Ti/Tv - Rare SNPs | 2.42 | 2.46 | 2.35 | 2.3 | 2.52 | 2.61 | 0.5 |

**Figure 5:**
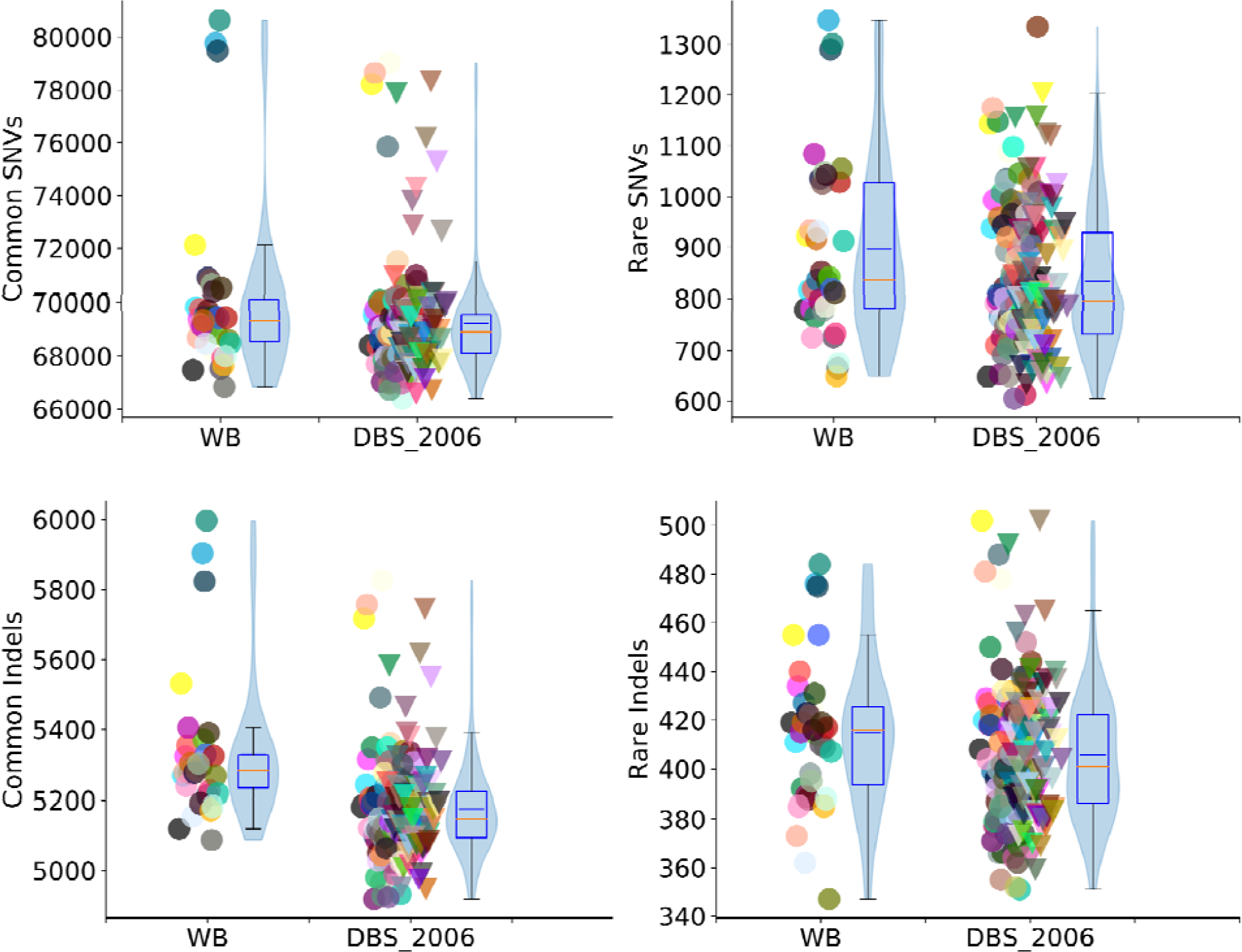
Hiqh quality variant counts from WB and DBS_2006 data. Individual sample values are plotted and adjacent box plots display the median (red) and interquartile ranges for the dataset, whiskers extend to the last data point within 1.5 times the interquartile range. Violin plots superimposed on the box plots show the data density and mean value (blue). Common variants have a frequency ≥0.001 and rare variants < 0.001 in 1000 genomes data (phase3). High quality is defined as marked PASS by VQSR and GQ≥30. All high quality variant counts were comparable except for common indels which were slightly lower in the dried blood set.

### Hiqh quality potentially false variants

Variants that were called with high confidence and not found in 1000 Genomes, yet shared by more than 2 samples were likely to be false positives that could be caused by systematic DNA damage to some sites. Excessive counts of such likely false positive (LFP) variants in DBS compared to WB samples could indicate systematic sequencing errors or alterations induced by DNA damage. Here, we examined only those variants that were marked as high quality post filtering and were likely false positives in the WB and DBS_2006 samples. There were fewer LFP SNVs in the DBS_2006 samples (Table 4; Additional file1, Figure S3B) and similar counts of LFP indels compared to the WB set (Additional file1, Figure S3C).

### Transition/Transversion ratios

As a measure of the quality of the filtered variant set, we compared the Transition/Transversion ratios categorized on variant frequency in the DBS_2006 data with those observed in the WB set. The transition to transversion ratios were comparable between the WB and the DBS_2006 datasets for both common and rare SNVs (Table 4; Additional file1, Figures S3E and S3F) and in a similar range observed in 1000 Genomes phase 3 data for Nimblegen capture region (range - 2.36 to 2.50)

## Discussion

Several quality metrics demonstrated that DNA isolated from DBS stored for prolonged periods at −20° C under desiccation is suitable for WES and clinical interpretation. The percentages of unmapped reads and low-quality bases in DNA from our DBS samples were comparable to DNA from our control WB set. While a lower percentage of on-target high quality read pairs was observed in the DBS_2006 set because of higher duplicate reads and lower on-target reads, sufficient numbers of high quality read pairs were available in the DBS_2006 samples to provide a median coverage proportionate to the raw reads. Uniformity of coverage is a critical parameter in exome sequencing projects, and despite the lower median coverage, the DBS_2006 samples yielded comparable or better uniformity of coverage to exomes from WB. Similar trends were observed for poorly covered and well covered exons in a selection of genes of interest, boosting confidence in DBS exomes for clinical use. DBS stored appropriately for >30 years showed value ranges similar to the more recent ones (data not shown).

The inferred sex matched the stated sex in all DBS_2006 and WB samples, and mismatches with the reference genome were also comparable to those observed in WB. Further examination revealed similar patterns of individual nucleotide changes between the WB and DBS_2006 sets, with the exception of T>A and A>T, which were in higher frequency in the DBS_2006 set, although in the same ranges as observed in the WB set.

To assess the impact of these differences for downstream analyses, high quality SNV nucleotide change frequencies were compared between the WB and DBS_2006 samples (Figure 6). All nucleotide changes were comparable between the WB and DBS_2006 sets except for C>T, marginally higher in the DBS set (Additional file2, Table S3), but in the same overall range. A>T and T>A had a similar distribution with the WB set despite showing differences in the DNA damage comparisons. We also examined the DBS_1980 set for nucleotide misincorporations and noticed that most of the nucleotide changes were marginally higher than both the WB and DBS_2006 set although in similar ranges (Additional file 1, Figure S6; Additional file 2, Table S10a). This, however, did not impact the nucleotide changes in the high confidence variants, which were in the same ranges as WB and DBS_2006 (Additional file 1, Figure S7; Additional file 2, Table S10b). We speculate that the longer storage time of the DBS_1980 samples (35 years versus 0 - 8 years) could have caused cumulative degradation resulting in higher rates of nucleotide misincorporation.

**Figure 6:**
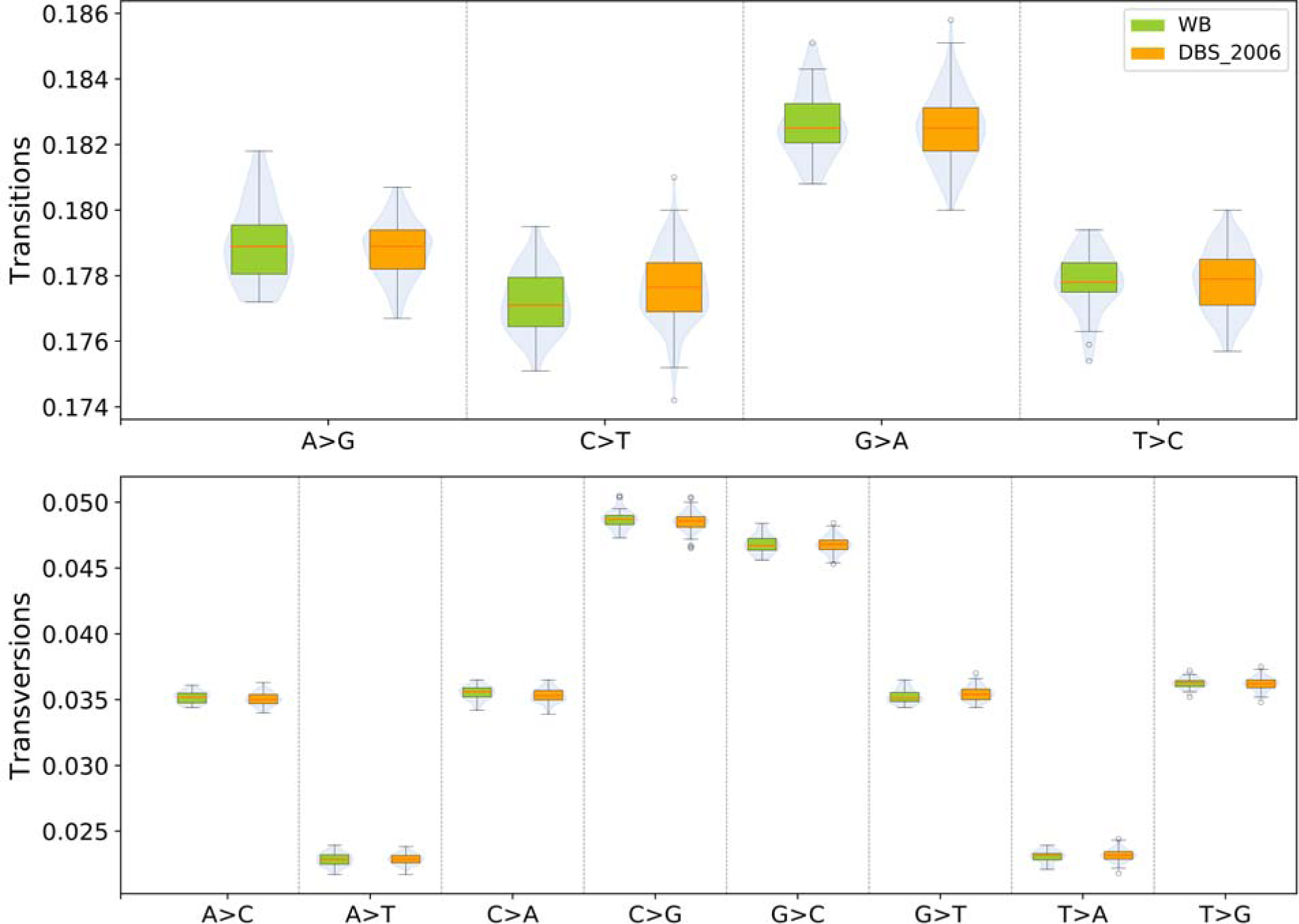
Frequencies for all single base changes in high quality. Box plots display the median and inter quartile ranges for the dataset, whiskers extend to the last data point within 1.5 times the interquartile range and outliers beyond this are marked with circles. High quality is defined as marked PASS by VQSR and GQ≥30. All changes were comparable in the 3 datasets except for the C>T fraction which was marginally higher in the DBS_2006 set.

High quality common and rare SNVs were comparable in the three sets. High quality common indels were slightly lower in the DBS_2006 set while rare indels were comparable in the both sets. We noted that the common indel counts before filtering were comparable between the WB and DBS_2006 sets (WB vs DBS_2006: P = 0.07; Additional file1, Figure S3D). Since coverage is also an important factor in determining variant counts, we performed a further comparison to rule out the effect of coverage on the indel counts. Samples with 50x to 80x median coverage were selected from the WB and the DBS_2006 sets and demonstrated comparable common and rare SNV and indel counts (Additional file 2, Table S9; Additional file1, Figure S4). False positive variant calls, which could result from base alterations due to DNA degradation, were identified with equal frequency for DBS_2006 and WB samples. Transition/Transversion ratios, a measure of quality of SNVs called, were also comparable in both sets.

## Conclusion

We explored several novel metrics to evaluate the quality of the DBS samples. These metrics are not restricted to the evaluation of DBS and have general applicability to exome sequencing data. Uniformity of coverage, which is independent of coverage depth, can be evaluated as a measure of confidence when yield of reads from sequencing is deemed insufficient. The coverage of the genes of interest in any study can be effectively reported visually with our gene coverage plots. Position-wise data across all exons is available at a glance with visual indicators to flag regions of poor coverage. The plots also highlight regions of exons not covered by the capture process. As an additional check of read quality, the sex of the sample can be inferred by our coverage, based sex identification approach. We have noted nonrandom distributions of nucleotide changes in our DNA damage metrics that are generalizable to other studies. Metrics such as confident sites and likely false positive variants give additional confidence to standard variant assessment practices.

In summary, our results indicate that DNA extracted from dried blood spots generally yields WES of comparable quality to DNA isolated from whole blood. The initial filter criteria that we applied to select samples for the study (Methods, Sample section), proved sufficient to short-list DBS samples for further analysis. Adequate quality coverage of the capture was achieved. Small differences in frequencies of nucleotide misincorporations did not impact the accuracy of variant calling. For DBS samples that have been archived for long periods, a preliminary additional check for DNA damage is suggested to ensure that the ranges are similar to whole blood samples. Thus, DBS can be used to sequence exomes suitable for clinical interpretation.

## Methods

### Samples

This study uses the following datasets for our comparisons:

- 180 De-identified, residual DBS samples (DBS_2006) that were collected and archived between 2006 and 2013 at −20°C with desiccant. The de-identified DBS were obtained from the CDPH California Genetic Disease Screening Program (GDSP) under an approved IRB protocol. The DNA was extracted and sequenced in 2015 for the samples.
- 35 de-identified WB samples (WB) were collected, extracted and sequenced between 2013 and 2014.

In addition, a smaller set of 8 anonymous DBS samples (DBS_1980) collected between 1980 and 1982 by CDPH were evaluated for the same metrics (Additional file 1; DBS_1980 metrics). These samples were archived at −20°C with desiccant.

This study was approved by the California Committee for the Protection of Human Subjects (project number 14-07-1650). DBS samples obtained from the GDSP as previously described [7] were used to prepare DNA for exome sequencing. DBS_1980 and DBS_2006 were archived by the GDSP and contained cases as well as false positives and controls for various metabolic disorders screened by MS/MS. WB samples were clinical exomes from children that were part of a separate study, but sequenced at the same facility.

Samples were selected to have a target capture coverage of at least 20x to ensure a minimum acceptable quality of exomes being compared. Contamination of the samples was estimated by the VerifyBamID tool [14] and samples with a freemix value >0.03 were excluded for the comparisons. A minimum fraction of 0.90 of high confident call sites (GQ≥30) across capture was ensured from the GVCF files. For the DBS_2006 set, this resulted in the selection of 180 out of 188 samples.

Comparisons were performed on various metrics between the WB and DBS sets in two ways – with just the WB vs the DBS_2006 set and also between the WB set vs DBS_1980 and DBS_2006 sets taken together.

The total number of reads were higher on an average in the WB samples compared to the DBS sets (WB vs DBS_2006: P value = 9×10^−5^; WB vs DBS_1980 + DBS_2006: P value = 9×10^−5^; Figure 1). There was more variability observed in the WB set and it should be noted that these were sequenced over multiple batches spread over 1.5 years.

### DNA Extraction, Target Enrichment and Sequencing

For DBS_1980 and DBS_2006 samples, sample preparation and sequencing were done as described [15]. Briefly, two 3-mm punches of a DBS filter were used for DNA extraction, libraries were prepared and pooled, and exon capture was performed by SeqCap EZ Human Exome Kit v3.0 (Nimblegen). Sequencing was done on a HiSeq 2500 (Illumina) sequencer to generate 100 bp paired end reads [16].

### Alignment and Variant Calling

Raw sequences were mapped to the reference genome (v37), using BWA mem algorithm (v0.7.10) [17]. Resulting SAM files were converted to binary format, sorted and lane merged using picard tools (v1.81) [18]. Duplicates were marked in the alignment files with Picard tools (v1.81). Next, realignment around known indels and base quality score recalibration were performed using GATK toolkit (v3.3) [19, 20]. Variants were called using the GATK Haplotype Caller function. The resulting VCF files were filtered using GATK VQSR function. Variant frequency and region annotations were added to the variant sets with Varant [21], an in-house annotation tool.

All computational analyses were performed using in-house python scripts unless otherwise mentioned.

### Reads and variants selection for metrics computation

Unless otherwise specified, only high quality read pairs (Additional file1; High quality reads pairs section) were used for analysis of the alignment and nucleotide misincorporation related metrics. Variant related statistics were reported for variants that were marked “PASS” and had a genotype quality ≥ 30 assigned by GATK Haplotype Caller.

### Metrics and statistics computation

All metrics were computed with python scripts and python libraries numpy and pysam, available for download from https://github.com/dbsexomes/DBSpaper.

Comparison of the various metrics between DBS and WB sets were done with Student T test (unpaired). Comparison was done two ways – i) WB vs DBS_2006 ii) WB vs DBS_1980 and DBS_2006 taken together. Since the parameter of comparison is source of DNA, DBS_1980 and DBS_2006 were not compared against each other.

## Additional files

Additional file1: Supplementary methods and figures (.docx)

Additional file2: Supplementary tables (.xlsx)

## Declarations

### Ethics approval and consent to participate

This study was approved by the California Committee for the Protection of Human Subjects (project number 14-07-1650). Residual, de-identified newborn dried blood spot samples were used to prepare DNA for exome sequencing. No sample or DNA remained after WES. If other researchers desire access to data or DBS, they would need to make a separate application to the CDPH (https://www.cdph.ca.gov/Programs/CFH/DGDS/Pages/cbp/default.aspx). California law precludes sharing specimens or uploading individual data derived from them into any genomic data repository.

### Availability of data and materials

Metrics generated from alignment and variant call files from whole blood and dried blood spot exome sequencing data are available in additional file 3.

### Authors contributions

US, RS, JMP and SEB conceptualized and designed the study; US generated the metrics from the exome sequencing data; US, ANA, KK and RS analyzed data. US, ANA, RS and SEB interpreted data. US, KK and RS created software. RS, JMP and SEB advised and provided suggestions; US wrote the first draft of the manuscript. RS, ANA, RJC, P-YK and JMP provided critical revisions. SEB was unable to review the manuscript for medical reasons. All authors (except SEB) approved the final version of the manuscript.

## Acknowledgements

The authors are grateful for expert technical discussions and computational assistance from Mark Kwale, John-Marc Chandonia, Dedeepya Vaka, Ajithavalli Chellappan, Navya Dabbiru, Brad Dispensa, Andrew Neumann, Yaqiong Wang, and Sadhna Rana. The biospecimens used in this study were obtained from the California Biobank Program, (SIS request number 496). The California Department of Public Health is not responsible for the results or conclusions drawn by the authors of this publication.

