## Supplemental methods and figures for "DNA from dried blood spots yields high quality sequences for exome analysis"

### Comparison of exome sequence quality from dried blood spot DNA with whole blood DNA

Supplementary methods and figures

#### Mapping quality metrics

**Unmapped Reads**
A large percentage of unmapped reads could indicate possible sequencing errors, or nucleotide misincorporations caused by damage in the DNA. We compared percentage of reads not mapped to any location on the reference genome by the aligner across the data sets. This metric was found to be slightly lower in the DBS_2006 compared to the WB samples (Additional file2, Table S1; Figure S1B). As a total percentage of the reads this was low in both sets (<1.8%).

**Low Quality Bases**
This is a measure of the quality of the sequences and impacts mapping quality and variant calls. Percentage of bases with base quality <20 were marginally higher in the DBS_2006 but in similar ranges to the WB set (Additional file2, Table S1).

**High Quality Read Pairs**
Properly paired reads, with both mates mapped to the same chromosome and in the correct orientation, non-duplicate and with mapping quality ≥20 were defined as high quality read pairs. These are the effective yield of the high quality reads from the sequencing. A significant decrease in high quality read pairs in the DBS samples relative to the WB samples could result in poor effective coverage of sequenced regions. The percentage of such on target high quality read pairs was lowest in the DBS_2006 set but not a function of the DNA source being comparable in the WB and DBS_1980 sets (Additional file2, Table S1; Figure S1C). On closer examination, we found that the DBS_2006 set had a higher percentage of duplicates and also a lower percentage of on target reads compared to the WB and the DBS_1980 samples (Figures S1D, S1E). Both of these impact the percentage of on target high quality read pairs.

**Read pairs mapped across different chromosomes**
We compared the fraction of read pairs that were mapped across different chromosomes by the aligner (BWA) using high quality read pairs described above. These were similar in both WB and DBS_2006 sets (Additional file2, Table S1) and an insignificant fraction of the total reads.

**Inferred Insert Size**
A significant percentage of reads with smaller than expected insert sizes can indicate fragmentation of the DNA due to damage. Median values for the inferred insert sizes as reported by the aligner (BWA) were calculated for each sample using high quality read pairs described previously. The DBS_2006 data could yield expected ranges of large insert sizes relative to the read length (Additional file2, Table S1; Figure S1F).

#### Variants – additional call sets

In addition to the combined call set of 223 samples consisting of WB, DBS_2006 and DBS_1980 sets (VCF1), additional call sets were generated:

VCF2: Variant call set with combined calls of 35 WB samples

VCF3: Variant call set with combined calls of 180 DBS_2006 samples.

Because of the low number of DBS_1980 samples, no combined call was generated for this data set.

This comparison was done in order to discount any influence data from the other sets might influence on the called variants. Both common and rare high quality SNVs were comparable in both the WB and DBS sets (Additional file2, Table S4; Figure S5). Rare indels were higher in the WB set as expected from the calling size effect of lower number of samples in WB (35 vs 180) which would result decreased filtering effectiveness of false positives. The Transition/Transversion ratios were higher for common SNVs were higher and lower for rare SNVs (Additional file2, Table S4, Figure S5). Likely false positive variants were also higher in the WB set. The WB samples from VCF2 also had significantly higher rare variants compared to the WB samples from the combined call set as a result of smaller call set size.

#### DBS_1980 metrics

The metrics for the 30 year old DBS samples were computed and compared with the WB set (Additional file2, Tables S5, S6, S7, S8). Since this was a small sample set (8), comparison was done by pooling this with the DBS_2006 samples and comparing the metrics based on DNA source. All the metrics showed similar trends as the DBS_2006 set with the DBS_1980 samples having good uniformity of coverage in spite of lower sequencing coverage, comparable amounts of DNA damage and similar variant ranges as the WB set. Nucleotide misincorporations were marginally higher for most of the base change types but did not impact the nucleotide change frequency in high quality SNVs or the SNV counts and quality (Figures S6 and S7; Tables S10a and S10b)

#### Code availability

The code used to generate the metrics from the alignment and call files is deposited in GitHub.

**Supplementary figures**

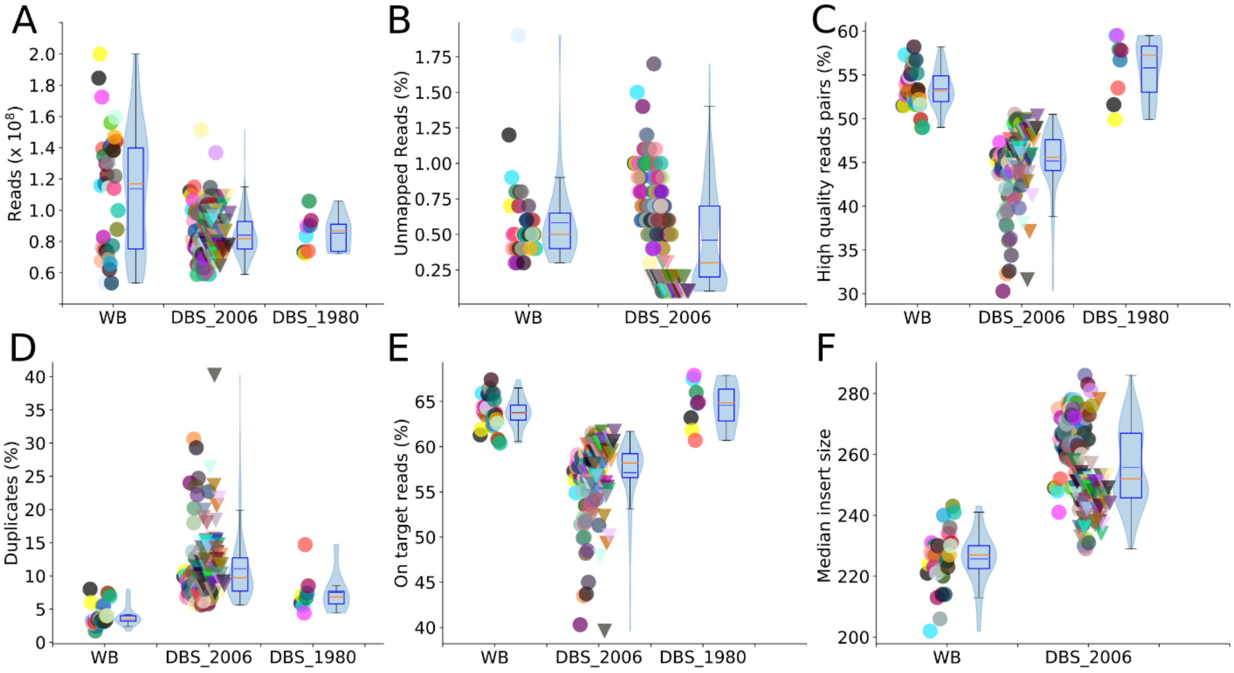


Figure S1: Mapping statistics in whole blood (WB), recent dried blood (DBS_2006) and old dried blood (DBS_2006) data sets.

Individual sample values are plotted and adjacent box plots display the median (red) and interquartile ranges for the dataset, whiskers extend to the last data point within 1.5 times the interquartile range. Violin plots superimposed on the box plots show the data density and mean value (blue). A: Number of reads in the 3 datasets. Many of the WB samples had a higher number of reads compared to the DBS_2006 dataset. The samples also showed more variance in terms of number of reads. B: Percentage unmapped reads. The percentage unmapped reads were slightly lower in the dried blood sets. All 3 datasets had low values (<1.8%). C: Percentage high quality read pairs. High quality read pairs percentage was lower in the DBS_2006 set. D: Percentage duplicate reads. Both dried blood sets had a higher duplicate percentage compared to the whole blood samples. E: Percentage on target reads. DBS_2006 had the lowest percentage of the 3 sets. F: Median insert size. DBS_2006 yielded expected insert sizes that were sufficiently large relative to the read length (100).

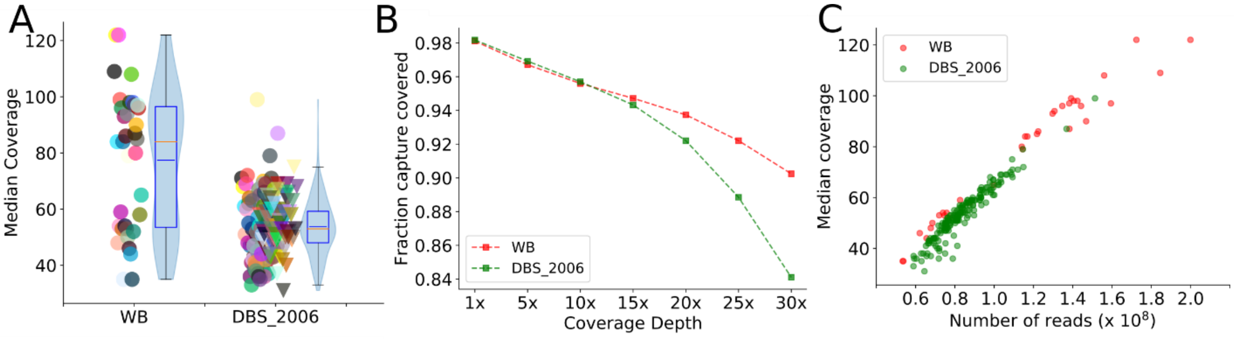


Figure S2: Median coverage and fraction of capture covered in whole blood (WB), recent dried blood (DBS_2006) and old dried blood (DBS_1980) data sets.

A: Median coverage was lower in the dried blood spot sets compared to the whole blood samples. Individual sample values are plotted and adjacent box plots display the median (red) and interquartile ranges for the dataset, whiskers extend to the last data point within 1.5 times the interquartile range. Violin plots superimposed on the box plots show the data density and mean value (blue). Fraction covered was comparable at 10x and fell post 20x in the DBS_2006 set compared to the whole blood data. B: Median fraction of capture regions covered (for each data set) at depths 1x to 30x for WB (red), DBS_2006 (green) and DBS_1980 (black). Fractions of capture covered was slightly higher in the dried blood sets at lower depths and fell above 15x depth. C: Median fraction of capture regions covered (for each data set) at depths 1x to 30x for WB (red), DBS_2006(green) and DBS_1980 (black). Fractions of capture covered was slightly higher in the dried blood sets at lower depths and fell above 15x depth.

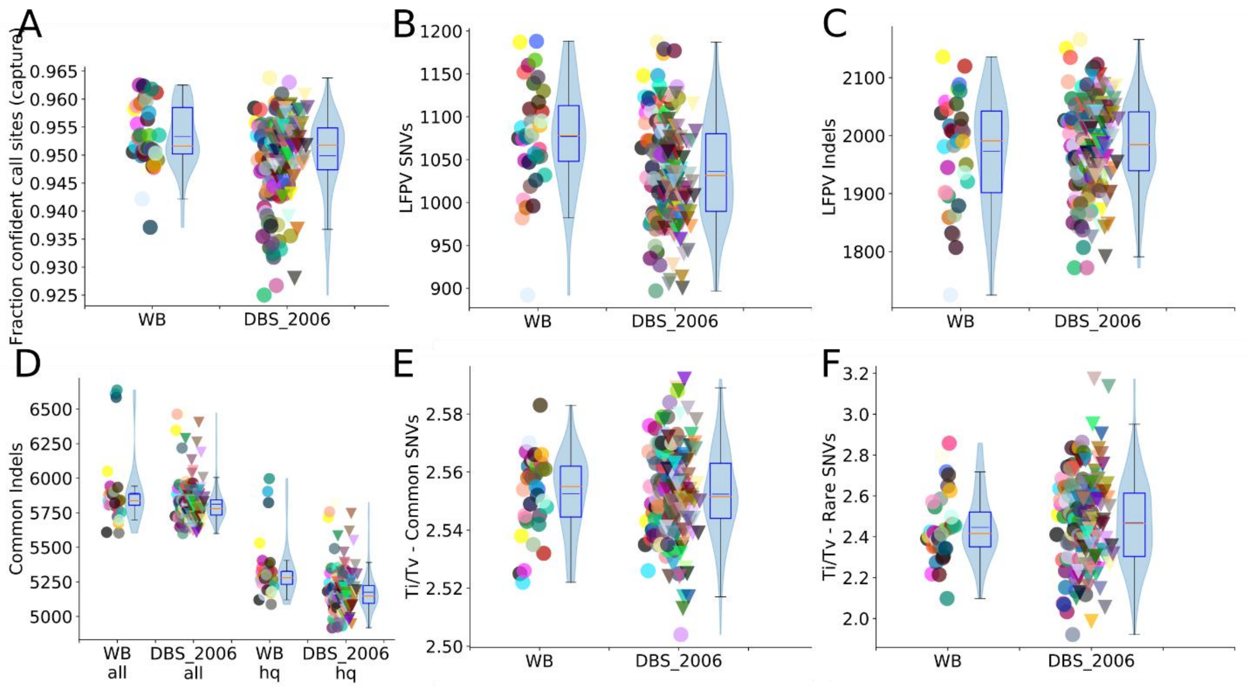


Figure S3: Variant statistics in whole blood (WB), recent dried blood (DBS_2006) and old dried blood (DBS_1980) data sets. In the figure, individual sample values are plotted and adjacent box plots display the median (red) and interquartile ranges for the dataset, whiskers extend to the last data point within 1.5 times the interquartile range. Violin plots superimposed on the box plots show the data density and mean value (blue).

A) Confident sites are those called either reference or variant in capture regions with GQ>=30 by GATK Haplotype Caller and calculated from the GVCF files. This was slightly lower in the DBS_2006 set compared to the WB and DBS_1980 set but the median value of the samples was > 0.95. B) & C) False positive SNVs and Indels are those that are marked PASS and with GQ>=30 and not present in 1000 Genomes database but shared in greater than 2 individuals. D) Common indels unfiltered and post filtering in the 3 datasets. Common is defined as >= 0.001 frequency in 1000 genomes data (phase3). High quality filters are marked PASS by VQSR and GQ>=30. Pre-filtering the indel counts are more comparable in the WB and DBS_2006 sets. Post filtering these are slightly lower in the DBS_2006 set. E) & F) Tranistion/Transversion ratios in hiqh quality common and rare SNVs. Common variants have a frequency >= 0.001 and rare variants < 0.001 in 1000 genomes data (phase3). High quality is defined as marked PASS by VQSR and GQ>=30.

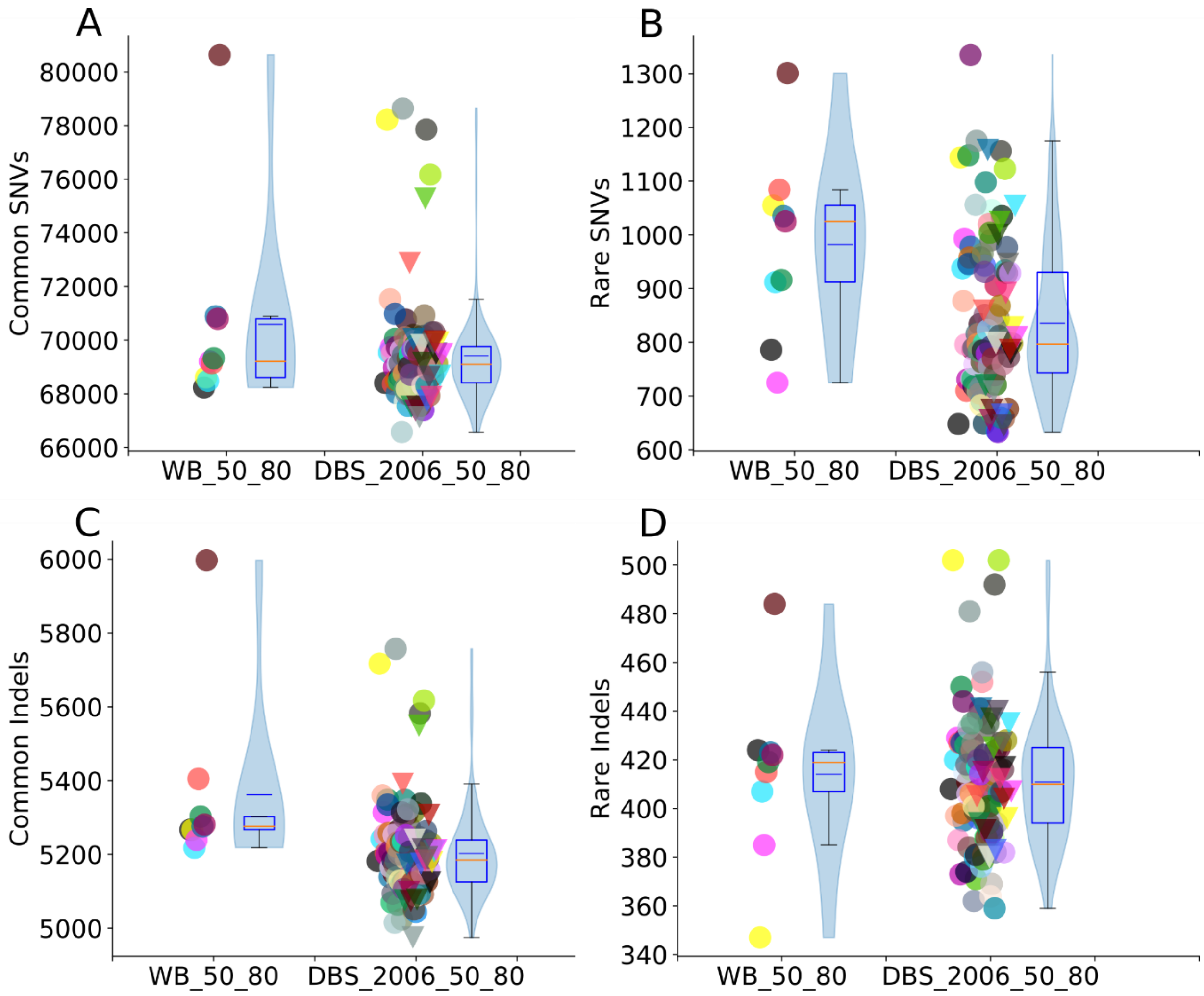


Figure S4: Hiqh quality variants in whole blood (WB) and recent dried blood (DBS_2006) with median coverage across capture between 50x and 80x. Individual sample values are plotted and adjacent box plots display the median (red) and interquartile ranges for the dataset, whiskers extend to the last data point within 1.5 times the interquartile range and outliers beyond this are marked with pluses. Violin plots super imposed on the box plots show the data density and mean value (blue). Common variants have a frequency >= 0.001 and rare variants < 0.001 in 1000 genomes data (phase3). High quality is defined as marked PASS by VQSR and GQ>=30. When the comparison was restricted to variants from samples with median coverage ranging between 50x and 80x, the counts of all high quality variants were comparable in the whole blood and dried blood data.

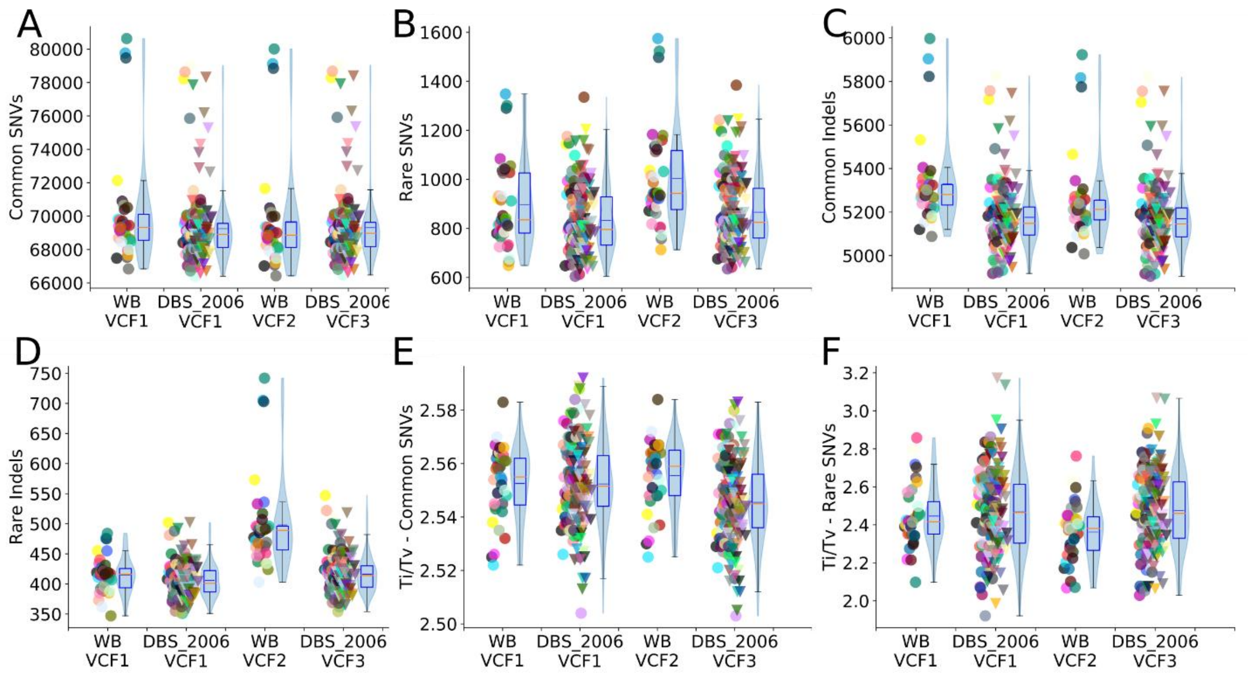


Figure S5: Variant statistics in whole blood (WB) and recent dried blood (DBS_2006) samples from VCF1, VCF2 and VCF3. Individual sample values are plotted and adjacent box plots display the median (red) and interquartile ranges for the dataset, whiskers extend to the last data point within 1.5 times the interquartile range. Violin plots superimposed on the box plots show the data density and mean value (blue). Common variants have a frequency >= 0.001 and rare variants < 0.001 in 1000 genomes data (phase3). High quality is defined as marked PASS by VQSR and GQ>=30.

A), B), C) and D) Hiqh quality variants. All high quality variants from VCF1 had comparable counts except for common indels which were slightly lower in the dried blood set. WB samples from VCF2 had a lower count of rare indels E) and F) Transitions to Transversions ratio (Ti/Tv) of common and rare SNVs.

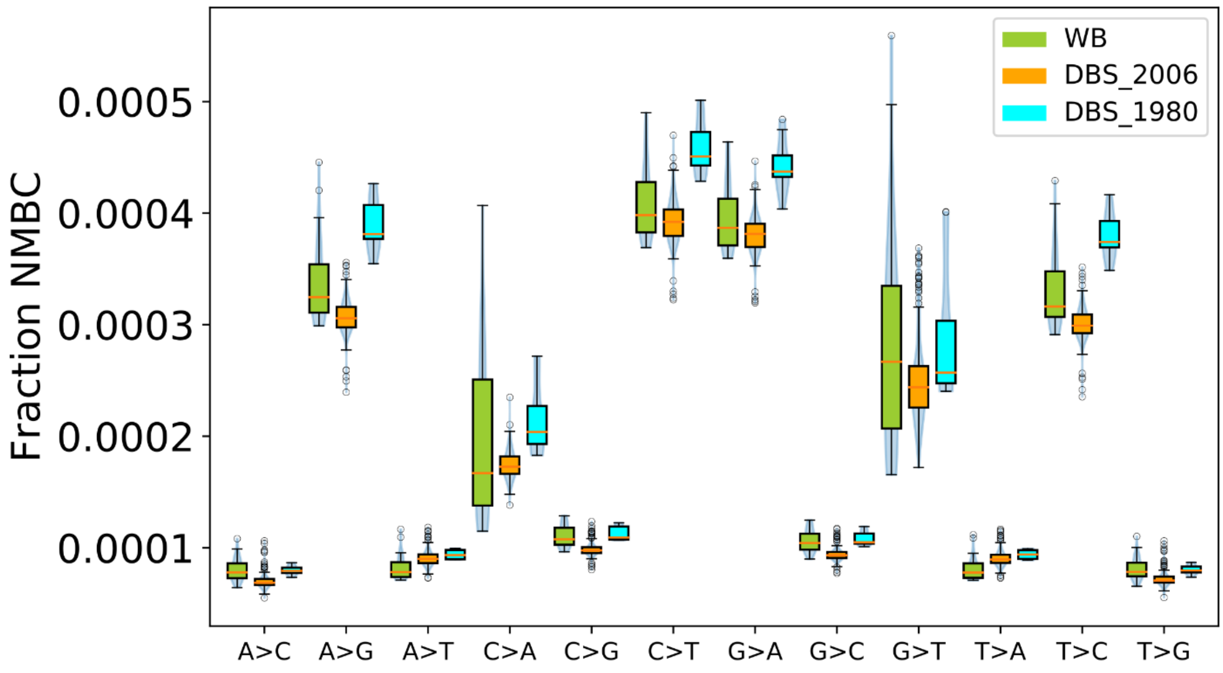


Figure S6: Fraction nucleotide misincorporations by base change (NMBC) for all single base changes. Box plots display the median and inter quartile ranges for the dataset, whiskers extend to the last data point within 1.5 times the interquartile range and outliers beyond this are shown as circles. Green: Whole blood (WB), Orange: Old dried blood spots (DBS_1980), Cyan: Recent dried blood spots (DBS_2006). Most of the nucleotide changes were marginally higher in the DBS_1980 set.

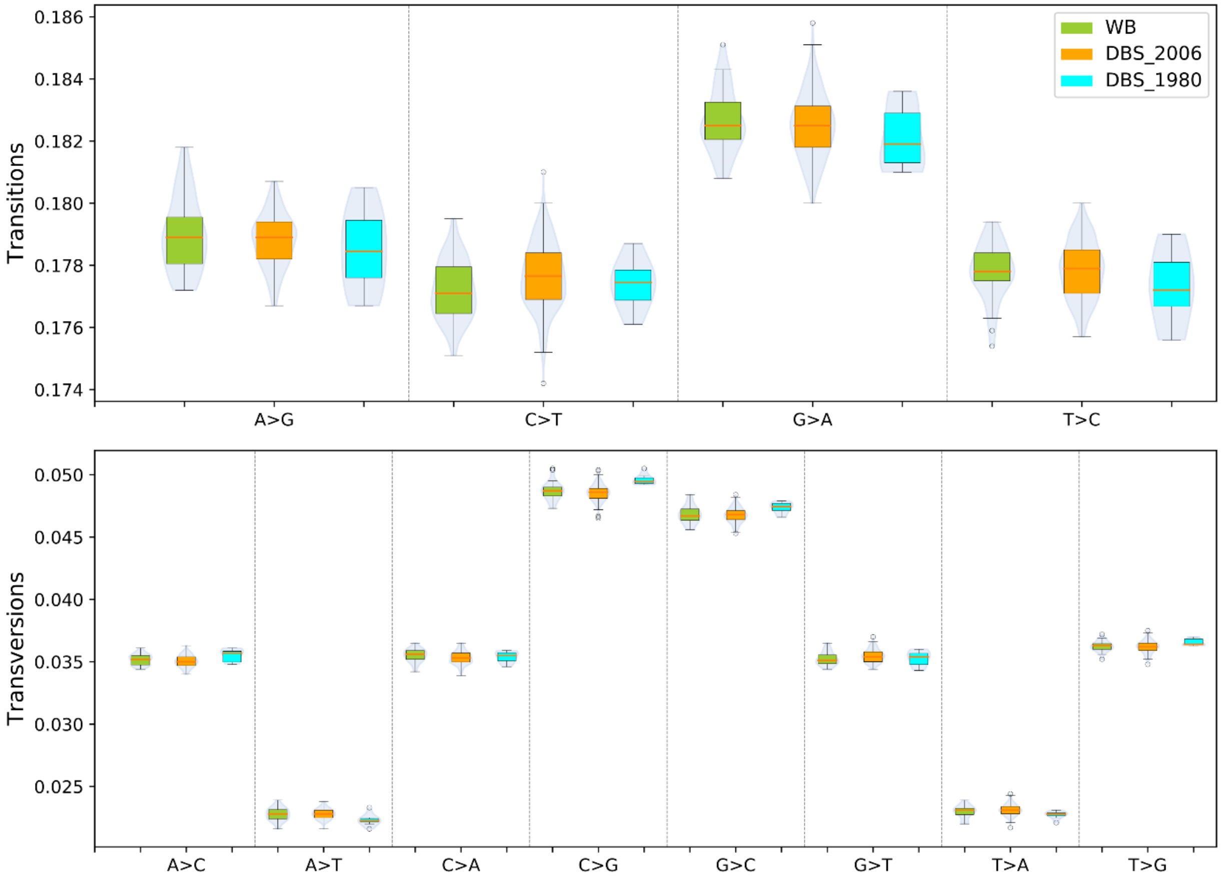


Figure S7: : Frequencies for all single base changes in high quality. Box plots display the median and inter quartile ranges for the dataset, whiskers extend to the last data point within 1.5 times the interquartile range and outliers beyond this are marked with circles. High quality is defined as marked PASS by VQSR and GQ>=30. All changes were comparable and in similar ranges in the 3 datasets.
